# MoveApps - a serverless no-code analysis platform for animal tracking data

**DOI:** 10.1101/2022.02.15.480513

**Authors:** Andrea Kölzsch, Sarah C. Davidson, Dominik Gauggel, Clemens Hahn, Julian Hirt, Roland Kays, Ilona Lang, Ashley Lohr, Benedict Russell, Anne K. Scharf, Gabriel Schneider, Candace M. Vinciguerra, Martin Wikelski, Kamran Safi

## Abstract

**Background:** Bio-logging and animal tracking datasets continuously grow in volume and complexity, documenting animal behaviour and ecology in unprecedented extent and detail, but greatly increasing the challenge of extracting knowledge from the data obtained. A large variety of analysis methods are being developed, many of which in effect are inaccessible to potential users, because they remain unpublished, depend on proprietary software or require significant coding skills.

**Results:** We developed MoveApps, an open analysis platform for animal tracking data, to make sophisticated analytical tools accessible to a global community of movement ecologists and wildlife managers. As part of the Movebank ecosystem, MoveApps allows users to design and share workflows composed of analysis modules (Apps) that access and analyse tracking data. Users browse Apps, build workflows, customise parameters, execute analyses and access results through an intuitive web-based interface.

Apps, coded in R or other programming languages, have been developed by the MoveApps team and can be contributed by anyone developing analysis code. They become available to all user of the platform. To allow long-term and cross-system reproducibility, Apps have public source code and are compiled and run in Docker containers that form the basis of a serverless cloud computing system. To support reproducible science and help contributors document and benefit from their efforts, workflows of Apps can be shared, published and archived with DOIs in the Movebank Data Repository.

The platform was beta launched in spring 2021 and currently contains 44 Apps that are used by 156 registered users. We illustrate its use through two workflows that (1) provide a daily report on active tag deployments and (2) segment and map migratory movements.

**Conclusions:** The MoveApps platform is meant to empower the community to supply, exchange and use analysis code in an intuitive environment that allows fast and traceable results and feedback. By bringing together analytical experts developing movement analysis methods and code with those in need of tools to explore, answer questions and inform decisions based on data they collect, we intend to increase the pace of knowledge generation and integration to match the huge growth rate in bio-logging data acquisition.

## Background

The growing field of bio-logging and animal tracking allows us to follow and document the movement behaviour and ecology of animals and species to an unprecedented extent and level of detail (Kays et al. 2015; Wilmers et al. 2015). However, as data volume and complexity have expanded, the extraction of knowledge has become increasingly challenging. The field of movement ecology has joined the big-data sciences: Tracking and bio-logging datasets comply with the “Four Vs Framework” (Volume, Variety, Veracity, Velocity) and their analysis “exceeds the capacity or capability of current or conventional methods and systems”(Farley et al. 2018).

For many users of bio-logging devices, the ability to fully exploit the information contained in tracking data increasingly lags behind the technological capacities (Holyoak et al. 2008). Some devices provide so much and such complex information that basic exploration of the data becomes a first major obstacle (Slingsby and van Loon 2016). As a result, experienced field biologists and wildlife managers must join forces with computational movement ecologists to process data appropriately in the quest to answer underlying ecological, management and conservation questions (Williams et al. 2020; Joo et al. 2020b). After collection, organisation and quality control, data are typically visually and analytically explored and processed in an iterative approach (Gupte et al. 2022). Following initial analysis, results often provide important insight leading to data re-analysis, data fusion (i.e. association with other ancillary information such as remote sensing data) or integration of additional data collected that were ignored in initial processing. This process results in new and often bespoke methodological workflows and analysis code (Reichman et al. 2011), but is tedious and not particularly sustainable or transparent (Lowndes et al. 2017) and requires accessory effort and investment to bring together the right combination of skills and interests in the research teams.

Ideally, these methods, workflows and analysis code compilations should be shared, compared, assessed and re-used or adapted across research groups and management agencies (Peng 2011). Indeed, many standard as well as novel analytic methods are being made available as open access code or functions in R-packages (Joo et al. 2020a; R-Core-Team 2021). R has become by far the most preferred software package for (movement) ecology, because it is open source and a large community contributes and maintains packages, continuously extending its scope and user community (Lai et al. 2019; Joo et al. 2020b). R-code and R-functions can allow efficient processing, exploration and robust analysis of datasets that cannot easily be accessed using software that has traditionally been used by field biologists (e.g. Excel or Google Earth). However, for some biologists, applied wildlife managers and those new to the discipline, the discovery, evaluation and use of this growing amount of code (Mislan et al. 2016) presents a major hurdle to being able to optimally benefit from state-of-the-art methods. Particularly for applied monitoring, conservation and management applications, it is of utmost importance that the information and insight gained from animal movement data can correctly and reliably inform decision makers, as well as support the possibility for near-real-time response when data are transmitted remotely from deployed tags.

The challenge of maximising the creation of knowledge from, and the beneficial use of, heterogeneous bio-logging data has been raised before, regarding the storage, standardisation and sharing of complex tracking datasets (Campbell et al. 2016; Sequeira et al. 2021). Online tracking databases have been established that allow researchers to stream, harmonise and store data from different types of tags, such as Movebank (movebank.org) (Kranstauber et al. 2011), Ocean Tracking Network (oceantrackingnetwork.org) and the EuroMammals family of databases (Urbano et al. 2010). These platforms perform vital steps to enable efficient analysis of tracking data, for example by standardising the coordinate reference system of location estimates and time zone and format of dates and times that define animal occurrences, and by providing shared data access protocols. In combination with appropriate metadata provided by data owners, long-term storage and exchange between researchers is made possible (Davidson et al. 2020). As sharing, publishing and combining data across groups and studies has become easier, so have collaborative projects and an interest in novel and accessible methods that can be applied to research, teaching, applied management and public engagement. The circle of ecologists participating in these databases has grown, increasing the taxonomic, geographic and temporal scope of harmonised data. At the same time, so has the community of developers contributing to the creation of new and innovative movement analysis methods, with potential to reach the crowd and citizen science community (Franzoni and Sauermann 2014).

One intrinsic complication of standardised and open software lies in the differences in development, maintenance, update and adaptation to novel computing infrastructure. Depending on requirements and preference, users may deploy new methods specific to particular working environments and operating systems that often are hard to combine. In addition, all operating systems need regular update and maintenance, and code and hardware degrade and become obsolete over time. Software packages require continuous code updates, and must often dynamically communicate with programmes and packages that also change regularly. The optimal utilisation of hardware resources at the currently highest performance levels requires additional maintenance and significant development effort. As a consequence, maintaining reproducibility of analysis code over long periods is very hard to achieve (Powers and Hampton 2019). Although code can be archived, including information about the used software, versions and settings, changing computing environments might make it near to impossible to execute code in future systems.

As a consequence of the above challenges and unexploited opportunities, the next step in improving the efficiency and benefits of analysis in movement ecology is, in our view, to foster more coordinated and inclusive cooperation between field ecologists, movement analysts and programmers. Such an effort could expand access to state-of-the-art methods and computing power, extend the community of experts that participate in analysis, support communication and exchange between those collecting data and those developing analysis methods, and secure reproducibility and scalability. Here, we introduce MoveApps (moveapps.org, Fig. 1), a web-based analysis-platform for animal movement data, developed with the aim to connect people who develop and drive the field of code analysis methods with people that use these tools for their newly collected datasets to answer research questions and inform decisions. The platform will make movement analysis methods more readily available and provide fast and tractable feedback, fostering communication across the range of skills and experience present in the research community. The platform enables data owners and analysts to work independently with opportunity for close exchange with each other. MoveApps is based on a serverless cloud computing system that is independent of changing infrastructure (Kearse et al. 2012; Perez et al. 2018), thus supporting long-term and flexible functionality of analysis code. The first beta version of MoveApps was released in February 2021.

**Figure 1.**
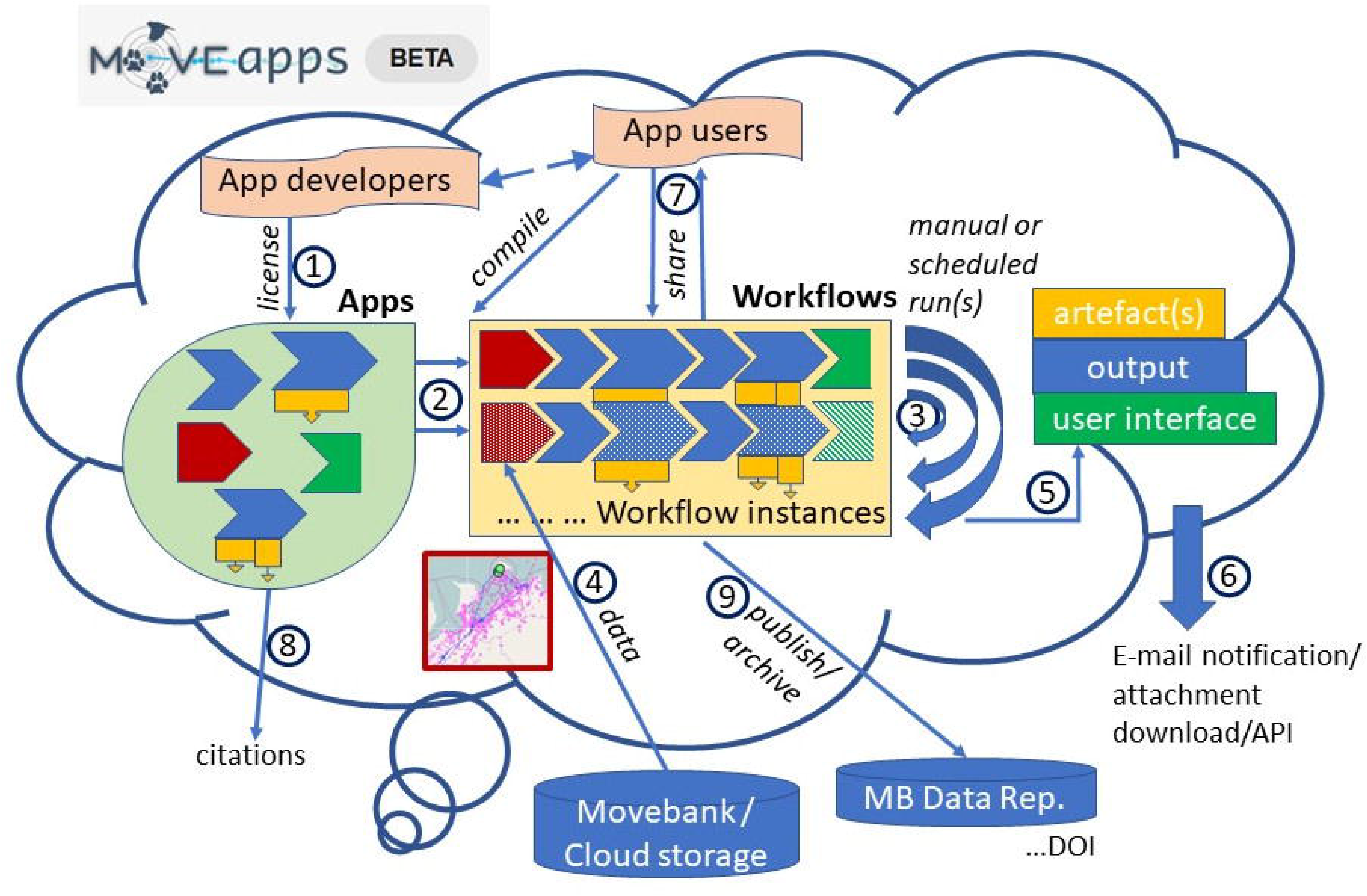
Schematic representation of the “cloud” computing MoveApps platform (beta version). (1) App developers provide Apps with defined input and output format under an open license. Apps can upload data (red), process data (blue), show results in an interactive user interface (green) and create products for download (yellow). (2) App users can combine those Apps to specific workflows to analyse their movement data. Workflows can consist of several workflow instances that can be (3) run manually or scheduled to analyse (4) tracking data. (5) The calculated results can be explored in a user interface or (6) downloaded as output and product files directly or via API. Notification E-mails can be sent of finished scheduled runs. (7) Workflows can be shared in the platform. (8) Citations for Apps are provided and (9) workflows can be published with a digital object identifier (DOI) and archived in the Movebank Data Repository. Registered MoveApps users can be App developers or App users (compiling workflows) or both.

## Implementation

### System requirements and design decisions

We designed MoveApps as a modular, open-source online platform that allows the secure use and exchange of interactive, user-developed analysis modules (Apps). Similar to other modular systems (e.g. Scratch (https://scratch.mit.edu), Node-RED (https://nodered.org)), the Apps can be linked and combined into data analysis workflows (Fig. 1). This modularity maximises flexibility and minimises each App’s complexity, likelihood for errors and development effort. Each individual App is meant to be a simple analysis building block that is defined by its input and output type. The analysis executed by an App is meant to be independent of a specific programming language or system structure. We specified MoveApps as a serverless platform (Perez et al. 2018) that runs on a cloud computing system, thus (1) operating independent of the users’ hardware, (2) providing reproducibility of workflows over a longer time, (3) supporting automated routines that can be applied to near-real-time data feeds and (4) allowing scalability to future high usage by distributed and scalable computing.

For compatibility with other systems and clouds, MoveApps was designed using widely used open-source tools and languages. Its platform background is programmed in Kotlin and Java (Ardito et al. 2020). For realising it as a serverless cloud computing system, we decided to implement Apps as containers instead of Virtual machines. Both provide virtual environments in which processes can run in isolation, but instead of emulating their own host operating system, containers share an underlying host (Cito et al. 2017). That makes them faster and requires less overhead, which is sensible for our platform of many different small Apps (coded by many different developers) working together. As underlying host, we use the open-source operating system Linux GNU. The two most widely accepted container systems for Linux are LXC and Docker (Bernstein 2014). In the light of distributed computing, we selected Docker for its better portability across machines (Boettiger 2015). Thus, each App runs as an independent module in its isolated Docker container with defined software language and package versions. This minimises cascading errors in overly variable, interconnected or interdependent sequences of Apps. The library of separately developed Apps in the form of Docker containers is automatically deployed, scaled and managed by Kubernetes (kubernetes.io), a widely used open-source container-orchestration system (Bernstein 2014). This system ensures that the Apps can interface and exchange their inputs and outputs in a safe and standardised way and supports scalability as the platform grows.

### App development

The base of our modular analysis platform are the Apps: each App is meant to be developed to independently perform one or a few main function(s) on the input dataset and then output its results for further handling to a subsequent App. Apps in development (Fig. 1.1) for the platform are managed in public Git repositories. Each repository contains the programme code for executing the App, a custom specification of the App and a documentation file adhering to our template. All functional development of the App code is done in the user’s typical compiler/editor and thoroughly tested before submission to MoveApps. In the first beta version of MoveApps, only R and R-shiny Apps are supported. We currently provide Software Development Kits with an initial R-Studio project that allows Apps under development to be locally run and perform as if they were launched on the MoveApps platform.

The required custom specification file (named appspec.json) can be compiled with the help of a settings editor that is provided on the MoveApps platform (moveapps.org/apps/settingseditor). This meta-information file must contain all parameter definitions, system dependencies, a selected license, language, keywords, author names and a link to the App documentation. Additional information that would be used during workflow publication can be specified, including references and funding sources. To improve metadata quality and interoperability with other services (Schneider et al. 2021), we have designed the structure and options to incorporate the DataCite metadata scheme (DataCite-Metadata-Working-Group 2021) and well-known identifiers, such as ORCID (https://orcid.org).

Each App requires a defined input and output type. The only input types currently supported are movement data in the “moveStack” format of the R-move-package (Kranstauber et al. 2020) or a specified .csv data frame that is internally transformed to a “moveStack”. Similarly, supported output types are “moveStack” data and, if the App can serve as a workflow endpoint, an interactive user interface (R-Shiny). The present limitation to “moveStack” ensures the proper use for movement analyses, but easy transformation to data frames and other formats in R allows future portability and the option to extend the range of interchangeable input data types. Apps can produce additional output “products” (previously called “artefacts”), which are files that can be downloaded from the MoveApps platform in various formats, such as .pdf or .csv. The dataset created as the output of each App can be downloaded in R format (.rds).

After initialisation of a new App in MoveApps, which includes the definition of the runtime environment, input and output data formats and provisioning of a link to the Git repository, a first App version must be created and submitted. Each submitted App version is checked by the MoveApps administrators for functionality and performant custom specifications. Upon passing this short review, the submitted App is wrapped in a Docker container and becomes available to all users on MoveApps. Improved App versions can be submitted at any time and become available to respective App users by notification of the possibility for update of Apps they used in existing workflows. All versions of an App are stored and can be reintegrated upon demand.

The use of an open-source language such as R, to which huge numbers of developers contribute, brings the challenge of interdependencies and possible inconsistencies as packages and the R environment are updated. Single Apps might cease to work properly. The limitation to a minimum number of necessary packages in an App will lower the probability of this to happen. However, due to the modular structure of MoveApps, a workflow can still run, if dysfunctional Apps are removed or replaced by similar but functional Apps, even if the output might differ. Thanks to the open source architecture and the metadata descriptions, the developers of malfunctioning Apps can be contacted by MoveApps users or administrators and the App can be updated, possibly in a joint effort via e.g. Git fork and pull requests.

### Empower the community to share and contribute

One major aim of MoveApps is to empower all members of the bio-logging and movement ecology community to easily contribute, use and benefit from the platform. Therefore, its dashboard is arranged in a user-friendly interface to intuitively browse and select data, Apps, App settings and options, and workflows by point-click-track. The users as well as App developers need not be familiar with or accommodate their work to the background infrastructure and can instead focus on their scientific or management questions and contents of the relevant data and Apps.

The MoveApps platform has been developed in the spirit of Open Science, sharing and joint improvement (Franzoni and Sauermann 2014; Nosek et al. 2015; Gewin 2016; Powers and Hampton 2019). While we are providing an initial offering of Apps and sample workflows, the bulk of development of Apps to the platform is meant to be taken over by a growing movement ecology community. A thorough user manual and tutorials (docs.moveapps.org) enable (i) App users to combine Apps and create workflows for analysis of their movement data and (ii) App developers to create and submit innovative Apps to the MoveApps platform for the community to discover and adopt. Over the past year, we have introduced the platform to potential users through workshops, conference presentations and meetings with dozens of government agencies, non-profit organizations and academic institutions, which have also served as an opportunity to identify pressing needs and prioritize functions to implement in the first phases of App development. We have introduced the platform to several user groups in conferences, workshops or by personal outreach. MoveApps is now integrated into multiple ongoing conservation-focused projects (e.g. https://ceg.osu.edu/animal-tracking-y2y), and additional workshops, user training sessions and hackathons are planned for 2022.

All submitted Apps must be provided under a selected open license for further use. We currently allow the choice between five widely used open (software) licenses: GNU General Public License, MIT License, GNU Affero General public License, 3-Clause BSD License and Creative Commons Attribution Share Alike (for more details see https://choosealicense.com/licenses/). Each of these options allows free use of the App by any App user in MoveApps as well as the copying of code for further use or archiving. This builds the basis for true reproducibility and iterative improvement of the data analysis process (Fidler et al. 2017; Powers and Hampton 2019).

The MoveApps Terms (moveapps.org/terms-of-use) clearly state that the user is responsible for evaluating the functionality and suitability of each App and workflow. MoveApps and App developers cannot be held responsible for errors or unexpected output in such a community supported open source project. However, App developers must not knowingly include malware and need to provide a current contact E-mail address. We foster an environment of active personal collaboration and productive exchange between App developers and with MoveApps to jointly improve the system and App usability. However, the containerised architecture of MoveApps allows for safe execution of code (because inputs and outputs are defined by the system) and provides the opportunity to withdraw Apps, for example if flaws are identifier that cannot be feasibly resolved, without breaking workflows permanently.

### Comparison with other movement analysis tools

Apart from the large list of R-packages that allow the analysis of movement data (Joo et al. 2020a), there are several standalone, specific software tools for movement ecology analyses ((Resheff et al. 2014; Calabrese et al. 2016; Dodge et al. 2021); see also www.movebank.org/cms/movebank-content/software). Compared to MoveApps, these tools require a local installation or data upload from the local computer, limiting repeatability across users/devices and usefulness for users without access to sufficient computing power. Furthermore, some existing applications are partly commercial, imposing licensing costs and subscription plans, and by that additionally increase the hurdles of interacting and analysing movement data. Monolithic standalone applications further suffer from potential obscurity of the actual functionality and the underlying algorithms of the implemented functions provided, and were often developed to meet a specific need, with limited support and intent to offer future growth in functionality or customisation to support user requests.

The one system in ecology that is somewhat comparable with MoveApps, even if not serverless, is R with R-Studio itself and shinyapps.io. R is open access and most people use it in a local install instance (server-based installations are possible). It allows the addition of packaged functions by the community, as well as exchange and collaboration via Git. However, R-Studio as frontend can only be used by coding, which is the hurdle that MoveApps attempts to overcome. Shinyapps.io is a commercial online platform that allows the deployment, sharing and use of R-Shiny Apps. One example is “ctmmweb” which allows easy calculation of various home range measures (Calabrese et al. 2021). Similar to above discussed standalone software tools, R-Shiny Apps tend to become bespoke and often monolithic tools, that are difficult to adapt and alter. With its modular container structure, multi-language design and open source availability of Apps, MoveApps overcomes those limitations. It allows flexible and parallel improvements and variations of Apps and workflows as a community service. We chose to prioritise integration of R and R-Shiny into MoveApps in part to encourage integration of functions from these existing popular analysis packages (Joo et al. 2020a) into the platform early on.

## Results

### Workflow compilation, use and scheduling

Within MoveApps, Apps can be combined into workflows (Fig. 1.2), which define an ordered set of steps to access, process and analyse data. The process of building workflows is simple and intuitive in the platform’s graphical user interface, where users can browse Apps, view details of an App’s developers, purpose and documentation and select chosen Apps to add to a workflow. The list of Apps is alphabetically ordered, includes a short description of each Every workflow starts with a core App that loads data into the system (Fig. 1.4). As MoveApps has been set up as a partner platform to the Movebank data base within the Movebank Ecosystem (Kays et al. 2022), it is most convenient to directly import animal movement data stored in Movebank using the “Movebank” App. This core App allows users to log into Movebank to browse and securely transfer data based on their user access permissions within the Movebank data base, which accommodates both public and controlled-access data, provides support to harmonize data to a shared format and vocabulary, and supports live data feeds (Kays et al. 2022). Relying on Movebank for input of data to MoveApps thus provides a secure method to share data between collaborators, allows users without access to data storage or a fast internet connection to input large data volumes, reduces problems in analysis caused by inconsistent or unknown data formats, and supports automated reporting procedures during data collection (see example workflows below). Alternatively, uploading data files (.rds or .csv) from a personal cloud folder (Dropbox, Google Drive) is supported. This option offers flexibility to prepare multi-study datasets prior to importing to MoveApps, as well as to support Apps that incorporate other local data sources as part of tracking data analysis. The data are then passed on to the next App in the appropriate format and processed accordingly.

**Figure 2.**
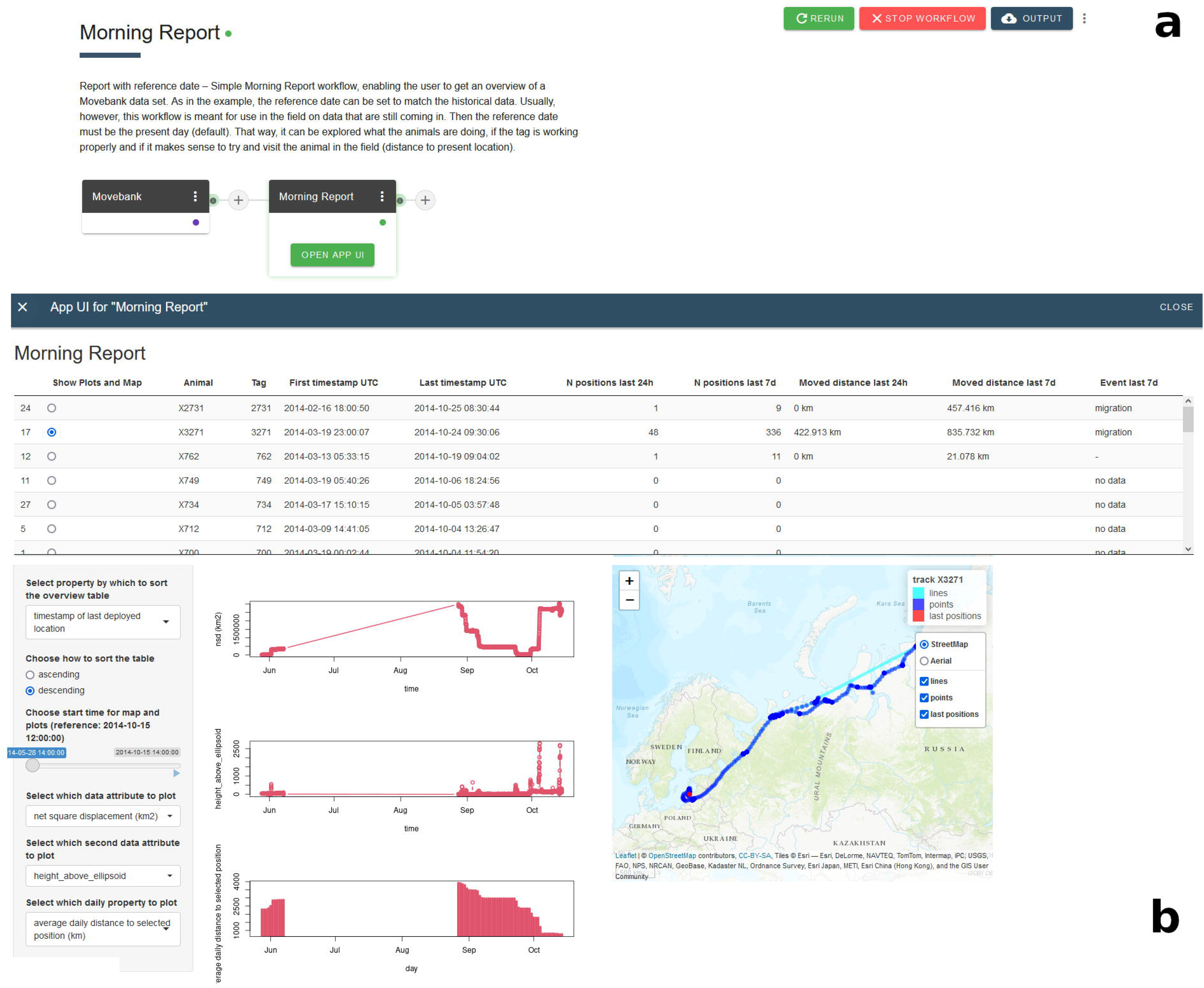
Example workflow “Morning Report”. Screenshots of the (a) workflow representation (order and names of combined Apps) and (b) workflow user interface output for an example dataset of greater white-fronted goose (*Anser a. albifrons*) tracks. Note that only tracks with data during 2014 were explored with the selected settings.

After data import, subsequent Apps can be added by selection from a list of all available Apps that accept the appropriate input and provide output in the required format. Input and output formats are filtered and matched automatically by the system. Once a workflow is compiled, it can be executed (Fig. 1.3). The user can follow the progress of each App in a workflow by the colour-indication of its state (idle, starting, working, post-processing or in error). Because MoveApps is cloud based, workflows run independently of the local machine and results from complex and time-intensive workflows can be checked after login at a later time. While the container structure of the workflow leads to somewhat longer runtimes in MoveApps than if the code was executed locally (see example workflows below), we consider this downside to be more than offset by the increased flexibility by users and other advantages of containers (see above). The workflow run can be stopped or re-started at any time. R-shiny Apps that invoke user interfaces can be opened after the App has finished and its results can be examined and users can interact with it according to the App’s programming features (Fig. 1.5).

**Figure 3.**
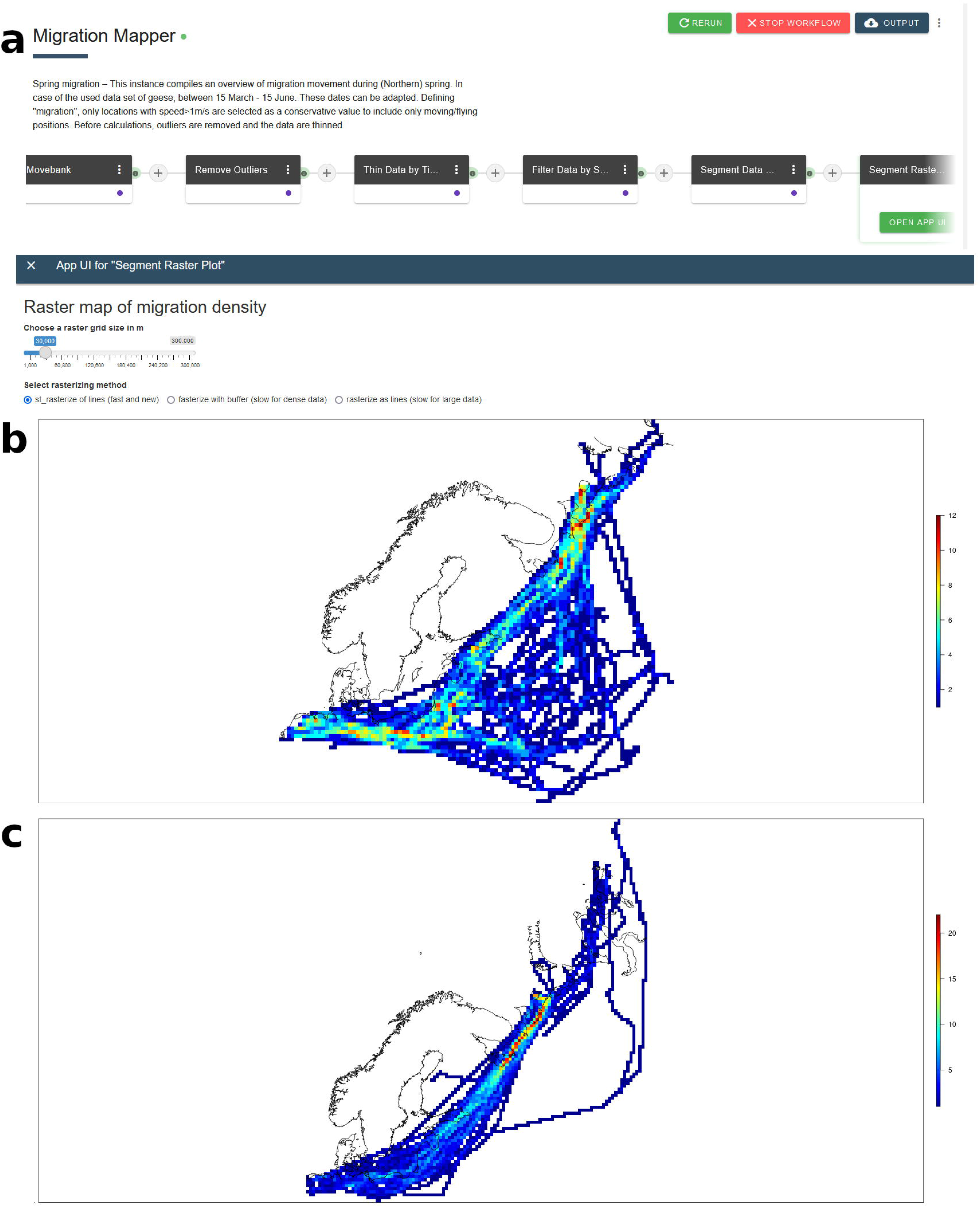
Example workflow “Migration Mapper”. Screenshots of the (a) representation (order and names of combined Apps) of the workflow instance “Spring migration”, (b) workflow user interface output for Spring migration and (c) Autumn migration of an example dataset of greater white-fronted goose (*Anser a. albifrons*) tracks. Note that tracks of all years are combined.

App details can be viewed at any time by opening the App menu. From this menu, user can change settings or access logs (process run, warning or error messages). Users can also “pin” a workflow at a certain App to retain the results of an App and all preceding Apps in the workflow. As a result, only subsequent Apps to the “pinned” App are re-executed when a workflow is re-started. The purpose is to avoid re-running e.g. initial data access and preparation steps that can be time-consuming with large datasets, thus providing ease of use when iteratively composing workflows and testing App settings. Each App that returns data also generates a short summary of the output data (e.g. time interval, number of animals and positions), which can be viewed easily at any time after the App has finished running. This allows the user to swiftly review App results, identify possible errors or unexpected results of the App, and better understand how each App relates to the workflow output. Finally, each workflow can be cloned into several workflow instances that analyse different datasets or are run using different user-specified parameter settings in one or more of its Apps. Managed by Kubernetes, this allows parallel execution for easy exploration of the influence of the workflow’s parameter space on the results. All workflows and their instances are saved in the user account for future reference.

Workflow instances can be started manually or scheduled to run automatically and without further interaction at fixed time intervals. This is especially useful when up-to-date information about tagged animals are required on a regular basis. Results of the scheduled runs can be accessed in the MoveApps platform or via a secure API (Fig. 1.6). Users have the option to request an E-mail notification after each scheduled run is completed, containing either a link to the MoveApps site for output access and download or including selected output files as attachments. The integration of alert notifications in the E-mail is e.g. possible with the “Email Alert” App.

### Share, cite and publish

For replication, collaboration or other joint work, it is possible to share workflows with other MoveApps users (Fig. 1.7). Workflows can be either shared publicly or with specific users. Recipients can load a shared workflow into their account’s dashboard and edit it there independently of the original workflow. It is possible to add two kinds of messages with shared workflows: (1) an open text field that allows the user to provide a brief description of the workflow and (2) a data source message which is by default filled with details of the dataset used by the original workflow creator. Thus, sensitive data are not transferred. Recipients of workflows must access the input data from their own accounts, which maintains the integrity of data sharing rights as managed by users in Movebank.

The importance of transparency and reproducibility based on open data and open code/methods has been repeatedly highlighted (Nosek et al. 2015; Fidler et al. 2017), especially if ecological applications are involved that can have important or controversial implications for science or management and are hard to impossible to replicate (Powers and Hampton 2019). Further, there is a need to ensure that researchers receive professional benefit and recognition for sharing code (Reichman et al. 2011). Therefore, MoveApps provides a citation for all Apps (Fig. 1.8) and offers the option to publish and acquire a digital object identifier (DOI) for workflows that are related to a published paper and dataset (Fig. 1.9).

To support reproducibility and comprehensive documentation of published analyses, the published workflows, their related Apps (including settings and source code) and metadata describing the operating system, libraries, packages and run-time versions used are archived in the Movebank Data Repository (Fig. 1.9). This is a free and well-established repository in the movement ecology community (Schneider et al. 2021; Kays et al. 2022) that provides persistent identifiers for future access and is accepted by scientific journals. The repository is developed in accordance with the FAIR (Wilkinson et al. 2016) and TRUST (Lin et al. 2020) data principles. For publication and archiving of workflows, users are required to provide a description of the workflow and each contained instance, the names of all contributors, funding sources and license type. Similar information for each App used in the workflow is extracted from their custom specification files. In combination with MoveApps’ serverless and modular structure, this archiving service helps to ensure the future reusability of code and replicability of published results, as well as the possibility to assess, modify and improve code and related analytical methods. For replication outside of MoveApps, archived workflows can be downloaded for use locally, and old R-environments and R-package versions can be accessed from the CRAN website. In addition, published workflows will be retained as publicly shared workflows on the MoveApps platform for easy discovery and reuse.

### Example workflows

We illustrate the use of MoveApps with two example workflows that address common analysis needs: using the “Morning Report” and the “Migration Mapper”, we analyse a published set of migration tracks of greater white-fronted geese (*Anser a. albifrons*; Movebank study: “Migration timing in white-fronted geese (data from Kölzsch et al. 2016)”, (Kölzsch et al. 2016a). These workflows were developed to showcase the use of the platform and discuss possible extensions to the beta version. The workflows have been made public on MoveApps to be used by all registered users and have been published in the Movebank Data Repository (Kölzsch and Wikelski 2021; Kölzsch et al. 2021).

The “Morning Report” workflow (Fig. 2a, doi:10.5441/001/1.h4c0p8bv, (Kölzsch and Wikelski 2021)) is made up of two Apps, the “Movebank” App and the “Morning Report” App, where the latter extracts an overview of a dataset with times of tag activity, plots of tag properties and a small interactive map. This is meant to be used for projects with active tags to explore tag performance, identify changes in behaviour and possibly find the animals in the field. Four Apps (called “Morning Report pdf Overview”, “Morning Report pdf Attribute Plots”, “Morning Report pdf Property Plots” and “Morning Report pdf Maps”) were recently developed, which can be combined into a workflow that provides “.pdf” product files of a time overview for all animals/tags, various data properties and track maps for download. These files can be taken into the field, sent by E-mail or accessed via API.

The user interface output of the workflow ((Fig. 2b) reveals that there were (at least) six different animals with available data during the past 5 months in the dataset. The time range, number of locations and distances moved are indicated. For the selected animal, we can see that from mid-June to the end of August, no data were available. After this period, autumn migration commenced and the large displacements and route are visible in the plots and map. To assess performance, we ran the workflow on both MoveApps and on a local installation of R-Studio. The workflow took 3:15 min to run on MoveApps, of which the longest part was taken up by loading the data (2:55 min). In comparison, on a local system R-Studio (IntelCore i7, 16GB RAM, Windows 10 64-bit), running the same code required 2:55 min in total, with 2:46 min for loading the data. Relative performance will vary based on the available processing power available to users outside of MoveApps.

The “Migration Mapper” workflow (Fig. 3a, doi:10.5441/001/1.7tq16jr8, (Kölzsch et al. 2021)) is a more complex workflow made up of six Apps that load data from Movebank, remove outliers, thin the data, filter by season, segment the data by speed and then plot the remaining locations as a density raster. The raster plot is provided as a user interface in which the user can change raster size for more detail vs. better visibility. The division of the workflow’s functionality into the many small Apps has notable advantages: Modular runs of independent Docker instances are more stable and run on less resources than one large, complex App. Furthermore, each App can be used in new workflows or can be replaced in the present workflow by different or more advanced App versions or Apps that have similar functionality.

The user interface outputs of the two different workflow instances show the routes of greater white-fronted geese during spring migration (Fig. 3b) and autumn migration (Fig. 3c). Densely travelled areas become visible by the heat map colours and indicate movement rather than resting, because only flight locations were selected using the “Segment Data by Speed” App. The maps confirm the known differences between the two migrations: During spring the geese fly in a wide front, using many different routes, whereas during autumn most of them use the coastal route which they pass quickly (Kölzsch et al. 2016b). The runtimes of the workflow for spring and autumn migration only differed minimally, each taking about 5:20 min on MoveApps, and 3:00 min on local R-Studio (see above).

## Conclusions

In a time of extreme growth of size and complexity of datasets (Wilmers et al. 2015; Joo et al. 2020b), we present the MoveApps platform as a unique tool to improve our ability to analyse movement data with the best methods in a comprehensible and efficient way. Our development showcases how movement ecology as a scientific community can be empowered to make analysis methods more accessible, in particular for to ecologists and wildlife managers. The platform offers opportunities for interactive participation by those less comfortable with command line programming, shared methods and collaboration across projects and agency jurisdictions, and management and research strategies that take advantage of dynamic monitoring and analysis of data as they are being collected.

Beyond its user-friendly interface, the MoveApps platform with searchable and citable Apps will help the community stay up to date with and explore the rapidly growing list of methods for movement data analysis (Joo et al. 2020a). Methods will become easily accessible as citable, reproducible and community-approved Apps and can be tested, compared and further improved by the community. In addition, the combination of Apps into workflows allows for an unprecedented ability to run more complex analyses. Hence, the MoveApps platform is intended to accelerate scientific work, discovery and collaboration between research groups and communities.

As a serverless cloud computing facility, MoveApps runs independently of soon outdated operation systems and can be scaled to the needs of the community (Talia 2013). It can provide computing power to researchers or communities that might not have such facilities at their home institutions or who work in the field. Presently, MoveApps is hosted on the cloud infrastructure of the Max Planck Society and is free for all users. As demand might increase in the future or the request for faster processing of workflows becomes critical, the use of Kubernetes orchestration in MoveApps allows distributed computing with the possibility to involve commercial partners like Amazon, Google, IBM, Microsoft, Baidu or institutional cloud computing resources for improved performance and scaling. This would come with the caveat that the running costs charged by the commercial service providers would need to be covered by the users justified by their need to analyse their data. We hope that this concept of flexible and integrative cloud-based analytics, deliberately designed to accommodate Open Science procedures can serve as a model for other research infrastructure applications in the future.

Finally, MoveApps provides a new way of making scientific research reproducible in all steps. Currently, scientific papers and datasets can be published with DOI, adding analysis methods complements this list and closes an often-encountered gap (Powers and Hampton 2019). Owing to its serverless structure, analysis methods and code in MoveApps can be permanently stored and are reproducible and openly accessible for use and improvement (Nosek et al. 2015). We believe this to be a necessary step to better promote Open Science and expect that our idea will be taken up by other research communities.

MoveApps launched its beta version in February 2021 and presently contains 44 functioning Apps that are used by 156 registered users. We invite the community to test it, provide feedback and contribute their own Apps and/or workflows. In the near future, we plan to provide more interfaces for communication between users and App developers, and include the capability to submit Apps in programming languages other than R. Based on community demands and as part of ongoing projects, Python and Matlab will be integrated next, but there are no technical restrictions to extending this range. The inclusion of additional data that are commonly used in analyses of animal tracking data, such as remote sensing information, will be further defined in the coming year with the addition of planned Apps that incorporate such sources. Additional App input and output formats will lead to different types of Apps which can be combined in various ways, leading to rapid growth and scalability of the system. To ensure an open invitation to participate and broad community input, we introduce the platform in its beta release, while the platform is available and offers basic functionalities, and while feedback can still drive the direction and priorities for future development. We encourage the community to contribute, exchange ideas and help define the future of MoveApps.

## Supporting information

Electronic supplementary material

## Availability and requirements

**Project name:** MoveApps

**Project home page:** https://www.moveapps.org Operating system(s): platform independent

**Programming language:** Kubernetes, Docker, Kotlin/Java, R Other requirements: none

**License:** General MoveApps Terms (https://moveapps.org/terms-of-use); selection of open software licenses for contributed Apps

**Any restrictions to use by non-academics:** none

## List of abbreviations

DOI: digital object identifier

## Declarations

## Consent for publication

Not applicable.

## Availability of data and materials

The example tracks of greater white-fronted geese are available from the Movebank Data Repository: https://doi.org/10.5441/001/1.31c2v92f (Kölzsch et al. 2016a) and can be accessed from the open Movebank study “Migration timing in white-fronted geese (data from Kölzsch et al. 2016)”.

The example workflows including R-code and system specifications are available from the Movebank Data Repository (https://doi.org/10.5441/001/1.h4c0p8bv (Kölzsch and Wikelski 2021) and https://doi.org/10.5441/001/1.7tq16jr8 (Kölzsch et al. 2021)) and are globally shared workflows in MoveApps (“Morning Report” and “Migration Mapper“).

## Conflict of interest

The authors declare that they have no competing interests.

## Funding

The initial development of MoveApps was funded by the Ministry for Science, Research and the Arts of Baden-Württemberg and the Knobloch Family Foundation.

## Authors’ contributions

KS, MW and AKS conceived and specified the idea for the platform. DG, CH, JH and BR set up, programmed and support the platform. AK and AKS programmed the Apps. AK coordinated the system development, wrote the user manual and supports users. SCD provided expertise of Movebank. AL, CMV and RK tested the platform and brought up improvements. GS, IL and SCD developed the publication and citation process. AK led the writing of the manuscript. All authors contributed critically to the drafts of the manuscript and gave final approval for publication.

## Acknowledgements

We are grateful to Michael Quetting for project coordination and to Babak Naimi for contribution to the very early conceptions of the MoveApps idea and its start. SCD acknowledges support from the NASA Ecological Forecasting Program Grant 80NSSC21K1182.

## Notes

### Competing Interest Statement

The authors have declared no competing interest.

