## Supplementary material for "MoveApps - a serverless no-code analysis platform for animal tracking data": Electronic supplementary material

**Abstract**

**Background**

Biologging and animal tracking datasets continuously grow in volume and complexity, documenting animal behaviour and ecology in unprecedented extent and detail, but greatly increasing the challenge of extracting knowledge from the data obtained. A large variety of analysis methods are being developed, many of which in effect are inaccessible to potential users due to remaining unpublished, being proprietary code and/or due to a lack of coding skills required to implement them.

**Results**

We developed MoveApps, an open analysis platform for animal tracking data, to make sophisticated analytical tools accessible to a global community of movement ecologists and wildlife managers. As part of the Movebank ecosystem, MoveApps allows users to design and share workflows composed of analysis modules (Apps) that access and analyse data stored in Movebank (or other data sources). Users browse Apps, build workflows, customise parameters, execute analyses and access results through an intuitive web-based interface.

Apps, coded in R or other programming languages, have been developed by the MoveApps team and can be contributed by anyone developing analysis code. To allow long-term and cross-system reproducibility, Apps have public source code and are compiled and run in Docker containers forming the basis of a serverless cloud computing system. To support reproducible science and help contributors document and benefit from their efforts, workflows of Apps can be shared, published and archived with DOIs in the Movebank Data Repository.

The platform was beta launched in spring 2021 and currently contains 32 Apps that are used by 75 registered users. We illustrate its use through two workflows that (1) provide a daily report on active tag deployments and (2) segment and map migratory movements.

**Conclusions**

The MoveApps platform is meant to empower the community to supply, exchange and use analysis code in an intuitive environment that allows fast and traceable feedback. By bringing together analytical experts developing movement analysis methods and code with those in need of tools to explore, answer questions and inform decisions based on data they collect, we intend to increase the pace of knowledge generation and integration to match the huge growth rate in biologging data acquisition.

**Keywords:** animal movement, movement ecology, biologging, method sharing, analysis code publication, reproducibility, self-growing, cloud infrastructure

### 1 Background

The growing field of biologging and animal tracking allows us to follow and document the movement behaviour and ecology of more and more animals and species to an unprecedented extent and detail (Kays, Crofoot, Jetz, and Wikelski, 2015; Wilmers et al., 2015). However, as volume and complexity of the data increase, the extraction of knowledge becomes increasingly challenging. The field of movement ecology has joined the big-data sciences: Tracking and biologging datasets comply with the "Four Vs Framework" (Volume, Variety, Veracity, Velocity) and their analysis "exceeds the capacity or capability of current or conventional methods and systems" (Farley, Dawson, Goring, and Williams, 2018).

For many users of biologging devices, the ability to fully exploit the information contained in tracking data increasingly lags behind the technological capacities. Some devices provide so much and so complex information that basic exploration of the data becomes a first major obstacle. As a result, experienced field biologists and computational movement ecologists must join forces to process data appropriately in the quest to answer underlying ecological, management and conservation questions (Williams et al., 2020). After collection, organisation and quality control, data are visually and analytically processed in an iterative approach. Following initial analysis, results often provide important insight leading to data re-analysis, data fusion (i.e. association with other ancillary information such as remote sensing data) or integration of additional data collected that were ignored in initial processing. This process results in new and often bespoke methodological workflows and analysis code (Reichman, Jones, and Schildhauer, 2011). This process is tedious and not particularly sustainable nor transparent (Lowndes et al., 2017) and requires accessory effort and investment to bring together the right combination of skills and interests in the research teams.

Ideally, these methods, workflows and analysis code compilations should be shared, compared, assessed and re-used or adapted across research groups or management agencies. Indeed, many standard as well as novel analytic methods are being made available as R-packages (Joo et al., 2020; R-Core-Team, 2021) or as open access code. Such code can allow efficient processing, exploration and robust analysis of datasets that cannot easily be accessed using software that has traditionally been used by field biologists (e.g. Excel or Google Earth). However, for some biologists, wildlife managers and those new to the discipline, the discovery, evaluation and use of this growing amount of code presents a major hurdle to being able to optimally benefit from these state-of-the-art methods. But particularly for applied monitoring, conservation and management applications, it is of utmost importance that the information and insight gained from animal movement data can correctly and reliably inform decision makers.

The challenge of maximising the creation of knowledge from, and the beneficial use of, heterogeneous biologging data has been raised before, regarding the storage, standardisation and sharing of complex tracking datasets (Campbell, Urbano, Davidson, Dettki, and Cagnacci, 2016; Sequeira et al., 2021). Online tracking databases have been established that allow researchers to stream, harmonise and store data from different types of tags, such as Movebank (movebank.org, Kranstauber et al., 2011), Ocean Tracking Network (oceantrackingnetwork.org) and the EuroMammals family of databases (Urbano et al., 2010). These platforms perform vital steps to enable efficient analysis of tracking data, for example by standardising the coordinate reference system of location estimates and time zone and format of dates and times that define animal occurrences. In combination with appropriate metadata provided by data owners, long-term storage and exchange between researchers is made possible (see e.g. Davidson et al., 2020). As sharing, publishing and combining data across groups and studies has become easier, so have collaborative projects and an interest in novel and accessible methods that can be applied to research, teaching, applied management and public engagement. Furthermore, not only is the circle of ecologists adding to the growing pool of data on an ever increasing range of animal taxa constantly growing, also the community of developers contributing to the creation of new and innovative movement analysis methods is growing and about to reach out into the crowd and citizen science community (Franzoni and Sauermann, 2014).

One intrinsic complication of standardised and open software lies in the differences in development, maintenance, update and adaptation to novel computing infrastructure. Depending on requirements and preference, users may deploy new methods specific to different working environments and operating systems that often are hard to combine. In addition, all operating systems are in need of regular update and maintenance. Software packages require continuous code updates and different programmes and packages that need additional work to communicate with each other in a dynamically and continuously changing setting. The optimal utilisation of hardware resources at the currently highest performance levels requires additional constant change and significant development effort. As a consequence, maintaining exact reproducibility of analysis code over long periods is very hard to achieve (Powers and Hampton, 2019), also as code and hardware degrade and become obsolete over time. Although code can be archived, including information about the used software, versions and settings, changing computing environments might make it impossible to execute code in future systems.

As a consequence of the above challenges and unexploited opportunities, the next step in promoting a more efficient movement ecology research is, in our view, to foster a more intimate and inclusive cooperation in the divided labour of data analysis between field ecologists, movement analysts and pure programmers. Thus, the community of contributors could be extended, maintaining communication and exchange intact, while securing reproducibility and scalability at lowest possible entrance barriers. With the MoveApps platform (moveapps.org, Fig. 1) we aim to bring people who develop and drive the field of code analysis methods closer together with people that use these tools for their newly collected datasets to answer research questions and inform decisions. The platform will make movement analysis methods more readily available and provide fast and tractable feedback, fostering communication between the range of skills present in the research community. The purpose is that the community of analysts and data owners are both enabled to work more independently yet in close exchange with each other. MoveApps is based on a serverless cloud computing system that is independent of changing infrastructure (Kearse et al., 2012), thus enabling long-term and flexible functionality of analysis code. The first beta version of MoveApps was released in February 2021.

#### 2 Implementation

MoveApps is designed as a modular online platform that allows the use and exchange of small, user-developed analysis modules (Apps). The Apps can be linked and combined to data analysis workflows (Fig. 1). Each individual App is meant to be a simple analysis building block that is defined by its input and output type. The analysis that is run inside an App is meant to be independent of a specific programming language or system structure. Each App performs one main function on the input dataset and outputs its results for further handling to a next App. Thus, the development effort for App developers is kept low. The modularity further minimises each App's complexity and likelihood for errors, and careful optimisation and adaptation for general usability is encouraged. During the development process, the App developer can ignore the background cloud infrastructure of MoveApps and need not to accommodate any software or hardware requirements.

The platform background of MoveApps is programmed in Kotlin (Java). Each App in this serverless cloud computing systems runs as an independent module in an isolated Docker container with its defined software language and package versions. This minimises cascading errors in overly variable, interconnected or interdependent sequences of Apps. The library of separately developed Apps in the form of Docker containers is automatically deployed, scaled and managed by Kubernetes (kubernetes.io), one of the most widely used container-orchestration systems. It ensures that the Apps can interface and exchange their inputs and outputs in a safe and standardised way and that scalability is retained with a growing MoveApps platform.

#### App development

Apps in development (Fig. 1(1)) for the platform are managed in a public Git repository. The repository contains the programme code for executing the App, a custom specification of the App and a documentation file. All functional development of the App code should be done in the user's typical compiler/editor and thoroughly tested before submission to MoveApps. In the first beta version of MoveApps, only R and R-shiny Apps are supported. We currently provide Software Development Kits with an initial R-Studio project that allows Apps under development to be locally run and perform as if they were launched on the MoveApps platform.

The required custom specification file (named appspec.json) can be compiled with the help of a settings editor that is provided on the MoveApps platform (moveapps.org/apps/settingseditor). This file must contain all parameter definitions, system dependencies, a selected license, language, keywords, author names and a link to the documentation. Additional information that would be used during workflow publication can be specified as well, including references and funding sources. To improve metadata quality and interoperability with other services (Schneider, Kölzsch, and Safi, 2021), we have oriented the structure and options towards the DataCite metadata scheme (DataCite-Metadata-Working-Group, 2021) and well-known identifiers, such as ORCID (https://orcid.org).

Each App requires a defined input and output type. The only input type currently supported is movement data in “moveStack” format of the R-move-package (Kranstauber, Smolla, and Scharf, 2020). Similarly, supported output types are ”moveStack” data or the App can serve as a workflow endpoint in an interactive user interface (R-Shiny). Apps can produce output ”artefacts”, which are files that can be downloaded from the MoveApps platform in various formats, such as .pdf or .csv. The dataset created as the output of each App can be downloaded in R format (.rds). The range of input and output formats as well as the types and kinds of artefacts will be extended in later release versions of MoveApps depending on the needs of the community and the platform development plans.

After initialisation of a new App in MoveApps, which includes the definition of the runtime environment, input and output data formats and provisioning of a link to the Git repository, a first App version has to be created and submitted. Each submitted App version is checked by the MoveApps administrators for functionality, performant custom specifications and possible issues regarding our Terms of use and security. Upon passing this review and quality control process, the submitted App is wrapped in a Docker container (see docker.com) and becomes available to all users on MoveApps. Improved App versions can be submitted at any time and become available to respective App users by notification of the possibility for update of Apps they used in existing workflows.

#### Empower a self-growing platform

One major aim of MoveApps is to empower all members of the biologging and movement ecology community to easily contribute, use and benefit from the platform. Therefore, its dashboard is arranged in a user-friendly interface with intuitive browsing and data selection, analysis modules (Apps) and workflows by point-click-track. The users as well as App developers need not to know about or accommodate their work to the background infrastructure and can focus on the scientific contents of their contributions.

The MoveApps platform has been developed in the spirit of open source science, sharing and joining in improvement (Franzoni and Sauermann, 2014; Gewin, 2016; Nosek et al., 2015; Powers and Hampton, 2019). While we have provided an initial offering of Apps and sample workflows, the bulk of development of Apps to the platform is meant to be taken over by a large and steadily growing movement ecology community. Thorough documentation and tutorials (docs.moveapps.org) are intended to enable (i) App users to combine Apps and create workflows for analysis of their movement data and (ii) App developers to create and submit innovative Apps to the MoveApps platform for the community to discover and adopt.

All submitted Apps must be provided under a selected open license for further use. We currently allow five widely used open (software) licenses: GNU General Public License, MIT License, GNU Affero General public License, 3-Clause BSD

License and Creative Commons Attribution Share Alike (for more details see https://choosealicense.com/licenses/). Each of these options allows free use of the App by any App user in MoveApps as well as the copying of code for further use or archiving. This builds the basis for true reproducibility and iterative improvement of the data analysis process (Fidler et al., 2017; Powers and Hampton, 2019).

The MoveApps Terms (moveapps.org/terms-of-use) clearly state that the user is responsible for evaluating the functionality and suitability of each App and workflow. MoveApps and App developers cannot be held responsible for errors or unexpected output. However, we foster an environment of active collaboration and productive exchange between App developers and with MoveApps to jointly improve the system and App usability.

### 3 Results

#### Workflow compilation, use and scheduling

Within MoveApps , existing Apps can be combined into workflows (Fig. 1(2)), which define an ordered set of steps to access, process and analyse data. The process of building workflows is simple and intuitive in our graphical user interface, where Apps can be easily browsed and selected. Each workflow is visually represented by connected containerised Apps, including access points to e.g. App details, settings or result overviews as well as buttons to initiate or stop workflow runs. Workflows can be saved, edited and run for specific use cases.

Every workflow starts with a core App that loads data into the system (Fig. 1(4)). As MoveApps has been set up as a partner platform to the Movebank data base within the Movebank Ecosystem (Kays et al. unpublished data), it is most convenient to directly import animal movement data stored in Movebank using the "Movebank" App. This core App allows users to log into Movebank to browse and securely transfer data based on their user access permissions within the Movebank data base, which accommodates both public and controlled-access data (Kranstauber et al., 2011). Alternatively, uploading data files from a personal cloud folder (Dropbox, Google Drive) is supported. The data are then passed on to the next App in the appropriate format and processed accordingly.

After data initialisation, further Apps can be added by selection from a list of all available Apps that accept the appropriate input and provide output in the required format. The list is alphabetically ordered, includes a short description of each App and is searchable by keywords. Once a workflow is compiled, it can be commenced (Fig. 1(3)). Since the site is cloud based, the workflow runs independently and results can be checked after login at a later time. The workflow run can be stopped or re-started at any time. R-shiny Apps that return user interfaces can be opened after the App has finished and its results can be examined and interacted with according to the App's programming features (Fig. 1(5)).

App details can be viewed at any time by opening the App menu. From this menu, settings can be changed or logs (process run, warning or error messages) accessed. Users can "pin" a workflow at a certain App to retain the results of an App and all Apps preceding it in the workflow. As a result, only subsequent Apps are re-executed when a workflow is re-started, the purpose being to avoid re-running e.g. initial data access and preparation steps that can be time-consuming with large datasets. Each App that returns data also generates a short summary of the output data (e.g. time interval, number of animals and positions), which can be viewed easily at any time after the App has finished running. This allows the user to swiftly check App results and pinpoint possible errors or unexpected results. Finally, each workflow may be cloned into several workflow instances that analyse different datasets or are run using different user-specified settings in one or more of its Apps.

Workflow instances can be started manually or scheduled to run automatically at fixed time intervals. This is especially useful when up-to-date information about tagged animals are required on a regular basis. Results of the scheduled runs can be accessed in the MoveApps platform. Users have the option to request an E-mail notification after each scheduled run completed, containing a link to the MoveApps site for output access and download (Fig. 1(6)).

#### Share, cite and publish

For replication, collaboration or other joint work, it is possible to share workflows with other MoveApps users (Fig. 1(7)). Workflows can be either shared publicly or with specific users. The recipients can load a shared workflow into their account's dashboard and edit it there independently of the original workflow. It is possible to add two kinds of messages with shared workflows: (i) a free text message that allows the user to provide a brief description of the workflow and (ii) a data source message which is by default filled with details of the dataset used by the original workflow creator. Thus, sensitive data is not transferred and the recipients of workflows have to access the data from their own accounts.

The importance of transparency and reproducibility based on open data and open code/methods has been repeatedly highlighted (Fidler et al., 2017; Nosek et al., 2015), especially if ecological applications are involved that can have important or controversial implications for science or management and are hard to impossible to replicate (Powers and Hampton, 2019). Therefore, MoveApps provides a citation for all Apps (Fig. 1(8)) and offers the acquisition of a DOI (digital object identifier) for workflows that are related to a published paper and dataset (Fig. 1(9)), addressing the challenge of ensuring that researchers receive professional benefit and recognition for sharing code (Reichman et al., 2011).

For permanent reproducibility, the published workflows and their related Apps (incl. settings and source code) will be archived in the Movebank Data Repository (Fig. 1(9)). This is a free and well-established repository in the movement ecology community (Schneider et al., 2021) that provides persistent identifiers for future access and is accepted by scientific journals. All stored files and information are treated under the FAIR (Wilkinson et al., 2016) and TRUST (Lin et al., 2020) data principles. Due to MoveApps' serverless and modular structure, the archived workflows can be replicated over a long time on the platform, independent of work environment updates. For possible replication outside of MoveApps, metadata of the used operating system, libraries, packages and runtime versions are collected. For publication and archiving of workflows, it is required to provide a description of the workflow and each contained instance, the names of all contributors, funding sources and license type. Similar information for each App used in the workflow is extracted from their custom specification files.

#### Example workflows

We illustrate the use of MoveApps with two example workflows that address common analysis needs: the "Morning Report" and the "Migration Mapper" with which we analyse a set of migration tracks of greater white-fronted geese (*Anser a. albifrons*; Movebank study: "Migration timing in white-fronted geese (data from Kölzsch et al. 2016)", Kölzsch, Kruckenberg, Glazov, Müskens, and

Wikelski, 2016). These workflows were developed to showcase the use of the platform and discuss possible extensions to the beta version. The workflows have been made public on MoveApps to be used by all registered users and have been published in the Movebank Data Repository (Kölzsch, Hirt, and Safi, 2021; Kölzsch and Wikelski, 2021).

The "Morning Report" workflow (Fig. 2a, doi:10.5441/001/1.h4c0p8bv, Kölzsch and Wikelski, 2021) is made up of two Apps, the Movebank App and the Morning Report App, where the latter extracts an overview of a dataset with times of tag activity, plots of tag properties and a small interactive map. This is meant to be used for projects with active tags to explore tag performance, identify changes in behaviour and possibly find the animals in the field. Four additional Apps were recently developed from the Morning Report App that provide ".pdf" artefact files of a time overview for all animals/tags, various data properties and track maps for download. These files can be taken into the field or sent by E-mail.

The user interface output of the workflow ((Fig. 2b) reveals that there were (at least) six different animals with available data during the past 5 months in the dataset. The time range, number of locations and distances moved are indicated. For the selected animal, we can see that from mid June to the end of August, no data were available. After this period, autumn migration commenced and the large displacements and route are visible in the plots and map.

The "Migration Mapper" workflow (Fig. 3a, doi:10.5441/001/1.7tq16jr8, Kölzsch et al., 2021) is a more complex workflow made up of six Apps that load data from Movebank, remove outliers, thin the data, filter by season, segment the data by speed and then plot the remaining locations as a raster. The raster plot is provided as a user interface in which the user can change raster size for more detail vs. better visibility. The division of the workflow's functionality into the many small Apps has notable advantages: Modular runs of independent Docker instances are more stable and run on less resources than one large, complex App. Furthermore, each App can be used in new workflows or can be replaced in the present workflow by different or more advanced App versions or Apps that have similar functionality.

The user interface outputs of the two different workflow instances show the routes of greater white-fronted geese during spring migration (Fig. 3b) and autumn migration (Fig. 3c). Densely travelled areas become visible by the heat map colours and indicate movement rather than resting, because only flight locations were selected by the segmentation by speed App. The maps confirm the known differences between the two migrations: During spring the geese fly in a wide front, using many different routes, whereas during autumn most use the coastal route that they pass quickly (Kölzsch, Müskens, et al., 2016).

### 4 Conclusions

In a time of extreme growth of size and complexity of datasets (Wilmers et al., 2015), we present the MoveApps platform as a tool to improve our ability to analyse movement data with the best methods in a comprehensible and efficient way. Our development showcases how movement ecology as a scientific community can be empowered to make analysis methods more accessible to ecologists who are less comfortable with command line programming.

MoveApps launched its beta version in February 2021 and presently contains 32 Apps that are used by 75 registered users. We invite the community to test it, provide feedback and contribute their own Apps and/or workflows. In the near future, we plan to provide more interfaces for exchange between users, set up an App wish list to allow guidance to App developers and include the capability to submit Apps in programming languages other than R. Based on demands and input from the community, Python is planned next, but there are no technical restrictions. The inclusion of additional data formats, like remote sensing information, for combined analyses is under discussion. Additional App input and output formats will lead to different types of Apps which can be combined in various ways, leading to rapid growth and scalability of the system. As of now, the basic functionalities exist and we encourage the community to take over, contribute and exchange ideas with us to guide future development of MoveApps.

### 5 Availability and requirements.

**Project name:** MoveApps

**Project home page:** https://www.moveapps.org

**Operating system(s):** platform independent

**Programming language:** Kubernetes/Docker, Kotlin (Java), R

**Other requirements:** none

**License:** General MoveApps Terms (https://moveapps.org/terms-of-use); selection of open software licenses for contributed Apps

**Any restrictions to use by non-academics:** none

### 6 List of abbreviations

DOI - digital object identifier

### 7 Declarations

#### Ethics approval and consent to participate

Not applicable.

#### Consent for publication

Not applicable.

#### Availability of data and materials

The example tracks of greater white-fronted geese are available from the Movebank Data Repository: https://doi.org/10.5441/001/1.31c2v92f (Kölzsch, Kruckenberg, et al., 2016) and can be accessed from the open Movebank study "Migration timing in white-fronted geese (data from Kölzsch et al. 2016)".

The example workflows including R-code and system specifications are available from the Movebank Data Repository ( https://doi.org/10.5441/001/1.h4c0p8bv (Kölzsch and Wikelski, 2021) and https://doi.org/10.5441/001/1.7tq16jr8 (Kölzsch et al., 2021)) and are globally shared workflows in MoveApps ("Morning Report" and "Migration Mapper")

#### Conflict of interest

The authors declare that they have no competing interests.

#### Funding

The initial development of MoveApps was funded by the Ministry for Science, Research and the Arts of Baden-Württemberg and the Knobloch Family Foundation.

#### Authors' contributions

KS, MW and AKS conceived and specified the idea for the platform. DG, CH, JH and BR set up, programmed and support the platform. AK coordinated the development, programmed the Apps and wrote the documentation. SCD provided expertise of Movebank. AL, CMV and RK tested the platform and brought up improvements. GS, IL and SCD developed the publication and citation process. AK led the writing of the manuscript. All authors contributed critically to the drafts and gave final approval for publication.

#### Acknowledgements

We are grateful to Michael Quetting for project coordination and to Babak Naimi for contribution to the very early conceptions of the MoveApps idea and its start.

### Figure captions

**Figure 1**. Schematic representation of the "cloud" computing MoveApps platform (beta version). (1) App developers provide Apps with defined input and output format under an open license. Apps can upload data (red), process data (blue), show results in an interactive user interface (green) and create artefacts for download (yellow). (2) App users can combine those Apps to specific workflows to analyse their movement data. Workflows can consist of several workflow instances that can be (3) run manually or scheduled to analyse (4) tracking data. (5) The calculated results can be explored in a user interface or (6) downloaded as output and artefacts files. Notification emails can be sent of finished scheduled runs. (7) Workflows can be shared in the platform. (8) Citations for Apps are provided and (9) workflows can be published with a digital object identifier (DOI) and archived in the Movebank Data Repository. Registered MoveApps users can be App developers or App users (compiling workflows) or both.

**Figure 2**. Example workflow "Morning Report". Screenshots of the (a) workflow representation (order and names of combined Apps) and (b) workflow user interface output for an example dataset of greater white-fronted goose (*Anser a. albifrons*) tracks. Note that only tracks with data during 2014 were explored with the selected settings.

**Figure 3**. Example workflow "Migration Mapper". Screenshots of the (a) representation (order and names of combined Apps) of the workflow instance "Spring migration", (b) workflow user interface output for Spring migration and (c) Autumn migration of an example dataset of greater white-fronted goose (*Anser a. albifrons*) tracks. Note that tracks of all years are combined.

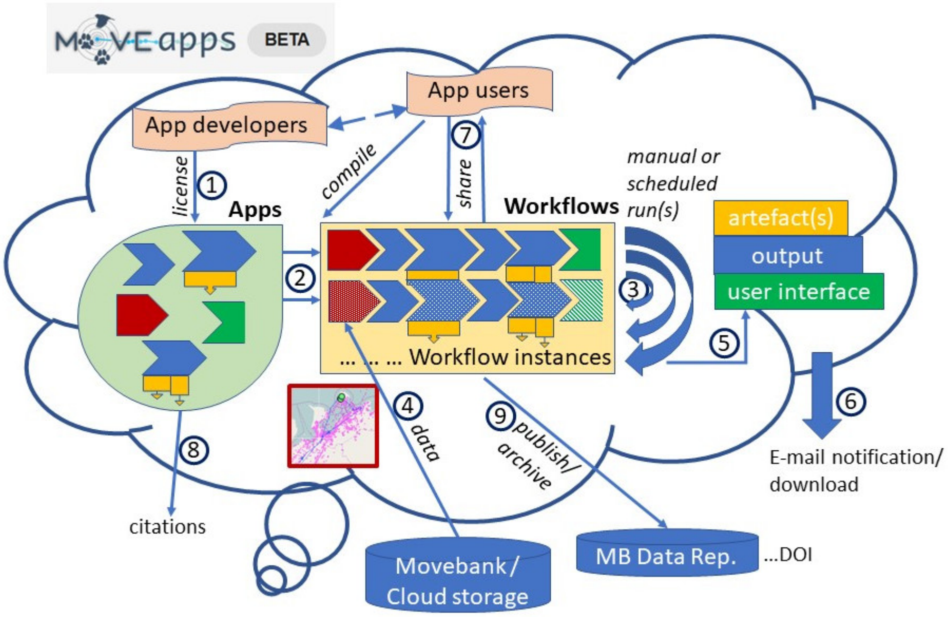


Figure 1: Schematic representation of the "cloud" computing MoveApps platform.

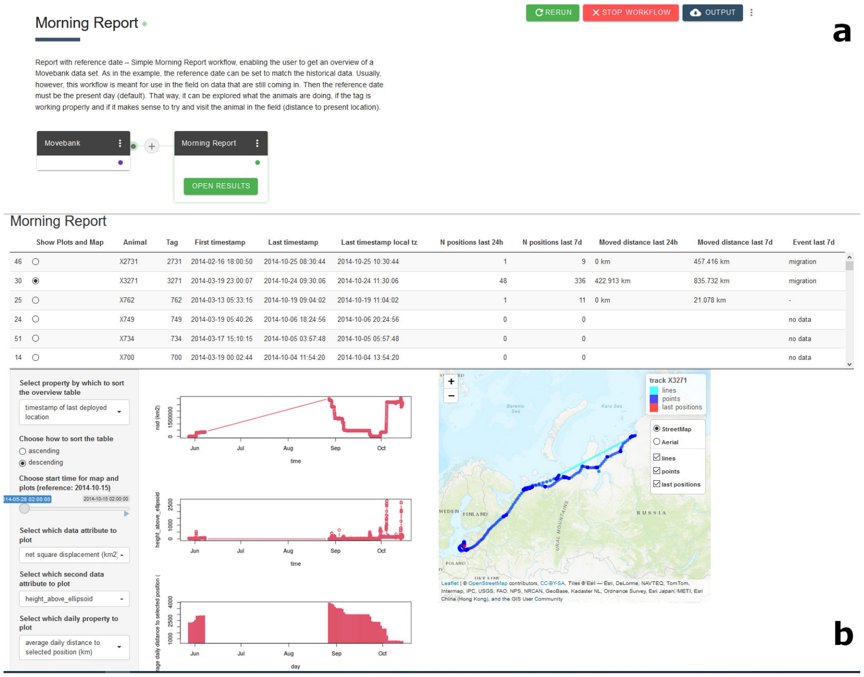


Figure 2: Example workflow "Morning Report".

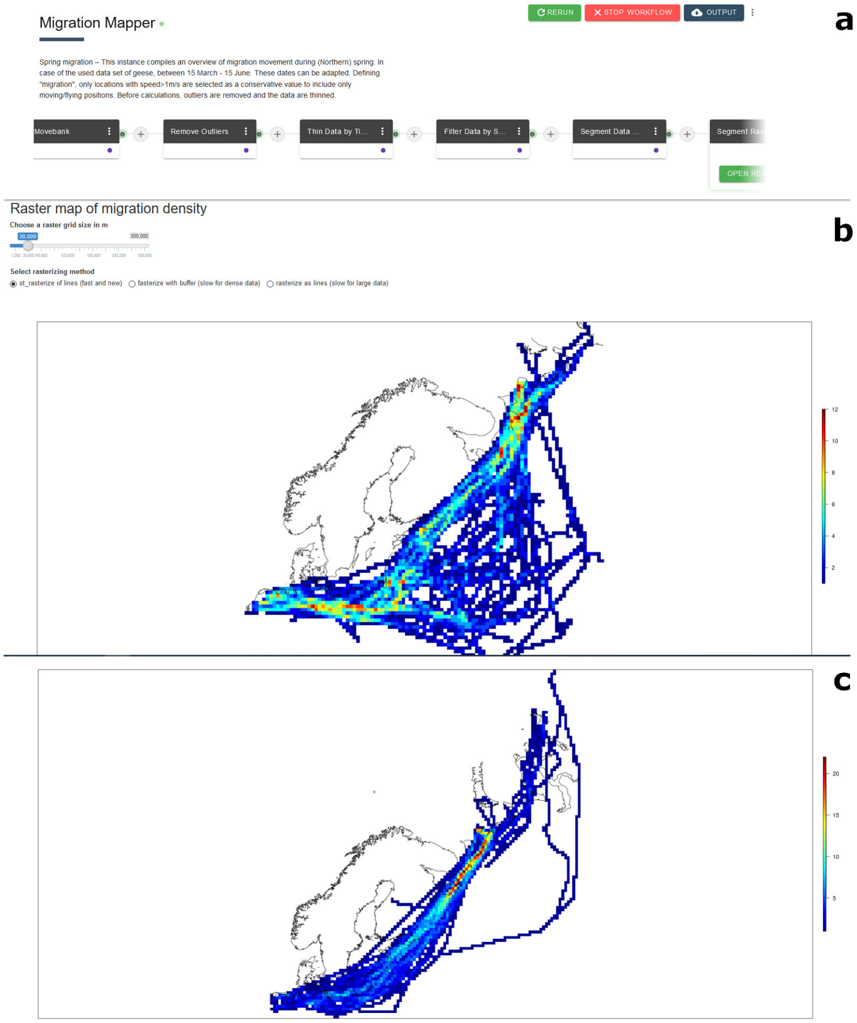


Figure 3: Example workflow "Migration Mapper".
